# Enhancement of neurophysiological signatures of working memory by combined yoga and tDCS

**DOI:** 10.1101/2023.09.18.558292

**Authors:** Omid Sefat, Mohammad Ali Salehinejad, Marlon Danilewitz, Reza Shalbaf, Fidel Vila-Rodriguez

**Author notes:** **Corresponding Author** Shared corresponding authors:, Reza Shalbaf, Fidel Vila-Rodriguez.

## Abstract

Transcranial direct current stimulation (tDCS) is a non-invasive neuromodulation technology that can modulate cortical excitability. Similarly, yoga has been found to influence neuronal activity and cognition. The aim of this study was to investigate the impact of combined yoga and tDCS on event-related potential (ERP) components during an N-Back working memory task. In a randomized, double-blind, cross-over design study, 22 healthy participants underwent a yoga/active tDCS session (2mA; 20min; anode on F3, cathode on F4) or a yoga/sham tDCS session on two different days. During the N-Back task, ERP components were obtained before and after each intervention. Results show that active tDCS plus yoga was associated with significant changes in the amplitude of the P200 component for the 2-Back in Fz and F3 channels and P300 for 3-Back in F3 and Pz electrodes. These results suggest that combining behavioral and electrical neuromodulation techniques may have the potential to enhance cognition and neurophysiological effects.

## Introduction

Yoga is a complex behavioral activity that combines aerobic physical activity with breathing and meditation-like exercises. Yoga has been shown to modulate brain activity with electroencephalography (EEG) and event-related potentials (ERP) [1,2,3]. Furthermore, regular yoga practice is associated with benefits on cardiorespiratory health, musculoskeletal disorders, and cognition [4]. In a recent meta-analysis yoga was shown to improve attention and processing speed, executive functions, and working memory [5]. Furthermore, correlates between neurophysiological and behavioral measures of working memory and executive function associated with yoga converged on modulation of frontal-parietal areas as a common underlying neuroanatomical underpinning [6,7,8].

Transcranial direct current stimulation (tDCS) is a non-invasive neuromodulation technique that delivers low-intensity direct current that elicits subthreshold neuromodulation of targeted brain tissue. Specifically, tDCS is associated with resting-state membrane potential changes that do not directly trigger action potentials but lead to meta-plasticity effects [9]. In healthy volunteers, tDCS is associated with positive effects on neurophysiological processes, and cognitive functions such as memory [9,10]. There is evidence that tDCS may enhance working memory by anodal stimulation of the dorsolateral prefrontal cortex (DLPFC) [11,12,13].

Working memory (WM) is the cognitive domain responsible for the storage of temporary data and information retrieval [14]. Working memory is the emergent product of interactions between long-term memory representations and fundamental processes, including attention, which are realized as reentrant loops between frontal and posterior cortical regions [15], where prefrontal cortex seems to have a primary generative role [16]. Working memory is a core cognitive function as it is associated with a wide range of complex cognitive abilities, such as learning, problem solving, reasoning and planning purposeful behaviors [17].

The N-Back task is a robust and one of the most frequently used neuropsychological test to probe WM. Concurrent use of EEG while performing N-back task to measure ERP is a powerful technique to interrogate the neurophysiological underpinnings of WM, particularly the components with latencies >100ms that are considered being more endogenous or cognitive such as the N200 (N2) or the P300 (P3) (i.e. these ERPs that are more sensitive to the subject’s interaction with the stimulus, as opposed to earlier ERPs which are more related to sensory components) [18].

Kesser and colleagues showed that tDCS (2 mA, 20 min) to the left DLPFC (anode on F3 and cathode above the right supraorbital region) in healthy individuals reduced error rate, and improved accuracy and reaction time in 2-Back. This was associated with an increase in the amplitude of the P200 and P300 components for the 2-Back task at the Fz channel [19]. Moreover, stimulation to the left DLPFC (anode on F3 and cathode on F4) using tDCS (15 min, 1mA and 2mA) in healthy individuals was associated with positive changes in the P300 component at Fz, a relationship that correlated with accuracy on the working memory test [20].

Only one prior study explored the combined effect of mindfulness and tDCS on working memory and its underlying neuropsychological mechanism. In this study, tDCS (2 mA, 30 min) administered on multiple sessions to the right frontal gyrus during mindfulness-based training sessions had a substantial effect on working memory and ERP features (P300 amplitude) in the stimulation area and Pz channel [21].

We examined the effects of yoga combined with tDCS over the prefrontal cortex on working memory ERP characteristics. We expected working memory indexed by the n-back task and its ERP/EEG neurophysiological aspects (P200 and P300 components) to change after combining yoga and tDCS stimulation on the left DLPFC. Specifically, we hypothesized that the neuromodulatory effect of tDCS and yoga would be noticeable in more demanding level of n-back task (i.e. 3-back rather 2-back) and neuroanatomically centered in frontal and parietal areas.

## Methods

### Participants

Twenty-two healthy volunteers were recruited. Two individuals withdrew consent before the first visit due to scheduling conflicts, and two participants’ EEG did not pass quality control (QC). Therefore, 18 individuals were selected for analysis, and reported here. Subjects were recruited by distributing brochures at yoga studios, at the University of British Columbia, local health food stores and, through online marketing, and by sending emails to university hospitals and yoga studio listserves. The criteria for inclusion were a) age between 18 and 35, and b) at least two years of consistent yoga practice. Exclusion criteria included a) pregnancy and b) persons afflicted by neurological disorders, brain damage, epilepsy, seizures, brain surgery, metal brain implants, or head trauma history. c) diagnosis of mental disorder as assessed by the Mini International Neuropsychiatric Interview (MINI), and d) lack of proficiency in the English language. According to the Declaration of Helsinki, the procedure was approved by the Ethics Committee at the University of British Columbia, and all participants provided written consent after being fully informed.

### Design

The study used a cross-over, randomized, double-blind design (Figure 1). Participants were randomly assigned to either yoga plus active-tDCS (a-tDCS) or yoga plus sham-tDCS (s-tDCS) on two different days with an interval of at least seven days between them. The tDCS was applied after yoga intervention. Sessions consistently occurred around the same time of the day to mitigate circadian variability [22]. Participants were randomly assigned using stratified randomization using the permuted block technique and a random number generator; the order of the a-tDCS vs s-tDCS sessions was counterbalanced. Demographic data were collected along with side effects. The combined yoga-tDCS sessions were flanked with collection of EEG before and after the yoga-tDCS intervention. Participants completed the N-Back task while EEG has been recorded in an Event-Related Potential paradigm.

**Figure 1:**
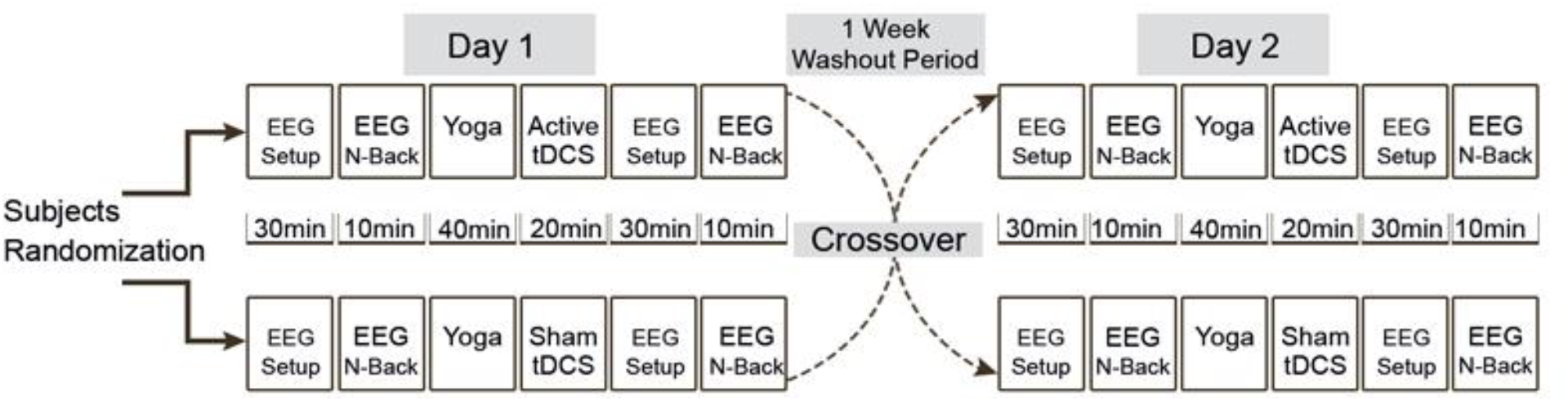
Methods and Study Design In this cross-over, randomized, double-blind trial, healthy participants got either yoga active-tDCS or yoga sham-tDCS. on two distinct days. Before and following each intervention, EEG/ERP data were collected during the N-Back task.

### Interventions

#### tDCS

A pair of saline-soaked surface sponge electrodes were used to supply direct current produced by an electrical stimulator (Newronika S.p.A., Milano, Italy) (surface of the electrodes 25 cm2). Anode electrode was placed over F3 (i.e., approximating the location of the left dorsolateral prefrontal cortex DLPFC) and cathodal electrode was on F4 (i.e., approximating the location of the right DLPFC) according to the 10/20 EEG system. The stimulation was administered in a quiet room where the participants sat comfortably while not engaging in any particular activity for 20 minutes at 2 mA current intensity (0.08 mA/cm2 current density). This montage was chosen based on research demonstrating its effect on cognition and executive functions, such as attentional control and working memory [12,23]. For the sham stimulation, the electrodes were placed at the same location but the stimulation current consisted of a 30-second ramp-up followed by a 30-second ramp-down, and no additional stimulation was delivered [24]. The tDCS researchers who investigated hypotheses and stimulation conditions were blind to stimulation conditions (i.e., active vs. sham). Each tDCS session was followed by a survey of adverse effects.

#### Yoga

The yoga intervention involved a 40-minute Hatha yoga class taught by a licensed instructor on a one-on-one basis. The posture sequence was developed by consulting with a professional yoga instructor and conducting a literature review [6].

#### N-Back

The N-Back letter task test was used to index working memory while recording EEG (Figure 2). The N-back task offers a fundamental assessment of working memory skills. Participants in the “N-back” task must identify stimuli presented n items earlier in chronological order, typically using letters. The difficulty of the cognitive challenge is modulated by the n-distance that has to be stored in memory, and it ranged between 1 to 3 back. Letter stimuli (20 capital consonants, excluding the letter ‘X’) were presented to participants in our investigation. Participants completed three consecutive runs of the test in increasing level of difficulty (i.e. 1-back, 2-back, and 3-back). Before each n-back session, participants were told through an instruction display whether that block would include an n-back (1,2 or 3) task. In each experimental session, one-third of the all 78 letter trials were targets. The stimulus presentation time was 500ms, the interstimulus interval (fixation time) was 2500 milliseconds. The stimuli were electronically delivered via E-Prime 2.0 software (Psychology Software Tools, Pittsburgh, PA).

**Figure 2:**
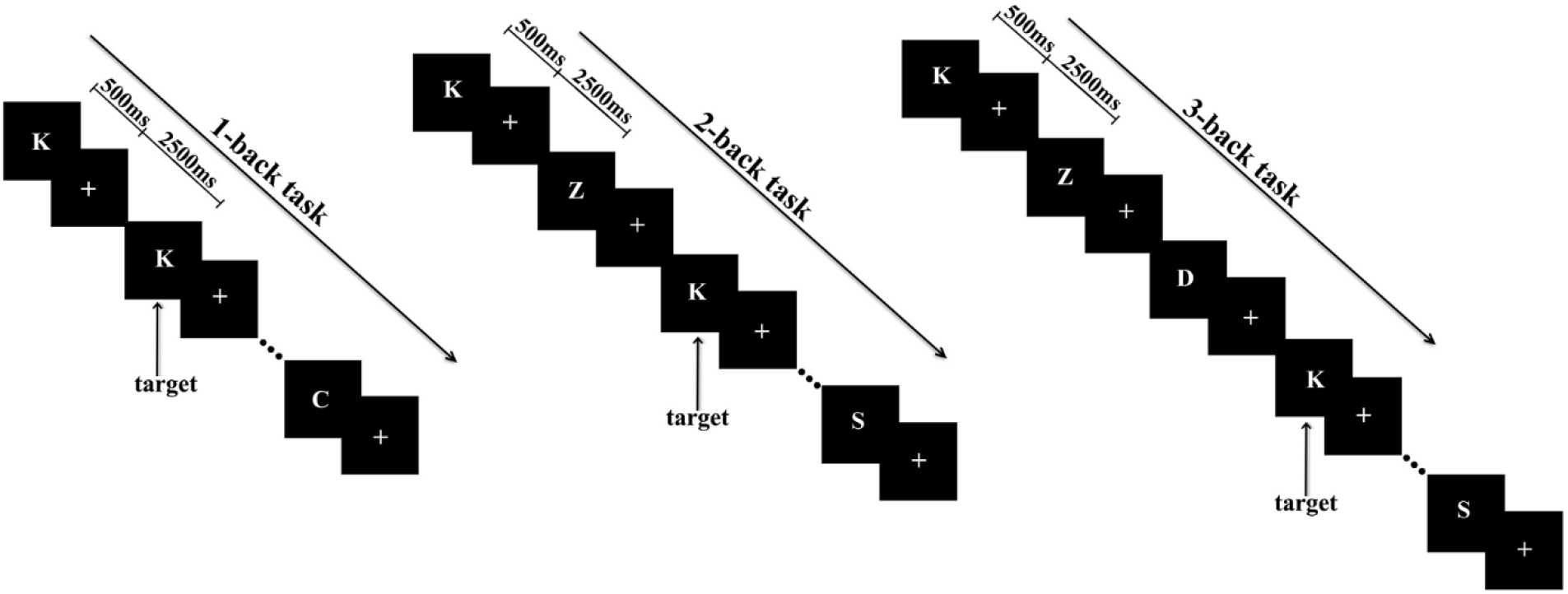
The format of the 1-Back, 2-Back, and 3-Back tests for working memory.

#### EEG

##### EEG Data Collection

BrainVision Recorder (BrainProducts®) was utilized to record EEG from 32 scalp channels (BrainCap, Brain Products®, GmbH, Gilching, Germany) placed in accordance with the 10-20 international standard. Cz was the location of the reference electrode, whereas Fz was the location of the ground electrode. By using Quik-Gel (Compumedics, USA) the electrodes attached to the head. The sampling frequency and band-pass filter were both set to 5000 Hz and 0-1000 Hz, respectively. Throughout the duration of the testing sessions, all impedances remained below 10 k. EEG information was collected in a quiet room prepared to minimize sources of electrostatic noise. Participants were requested to estimate their 24-hour intake of coffee, nicotine, sleep, alcohol, and over-the-counter medications before each session. Preparation of the subject for the EEG took approximately 30 minutes (cap setup). Except for TP9, TP10, VEOG, and HEOG channels, 28 of 32 electrodes were employed in the study (Supplementary materials). The frontal electrodes and stimulation site (Fz, F3, and F4), central (Cz) and parietal (Pz) were selected for analysis.

##### EEG signal preprocessing

In order to preprocess the EEG data and removing the artifacts, stages from Makoto’s preprocessing pipeline [25] were implemented in MATLAB 2020a using the EEGLAB toolbox 2020 (The MathWorks, Natick, MA) [26]. Specifically, data were resampled to 512Hz, high-pass filtered at 1 Hz, and then re-referenced to a mean reference. *CleanlineNoise* EEGLAB-Plugin was used for line noise elimination. Continuous data were cleaned using the EEGLAB plugin clean raw data, ASR (Artifact Subspace Reconstruction), A method that eliminates, low-frequency drifts, noisy and flatline channels, and brief bursts. Deleted electrodes were interpolated using spherical interpolation (Mean=1.2 with a range of (0,3)). Then, the entirety of the unprocessed data was visually examined to identify artifact-relevant parts In order to reduce non-brain artifacts in EEG data, Independent components (ICs) were obtained through adaptive mixture ICA (AMICA). We distinguished between brain ICs and other independent components (heart, eye, muscle, brain, etc.) using the ICLabel plugin in EEGLAB, with a probability of ‘brain’ label greater than 0.7.

##### ERP analysis

ERP analysis was conducted for the hits (correct responses to the target stimuli) and non-target correct rejections. The preprocessed EEG data were split into epochs between −250 and 750 milliseconds relative to stimulus onset. In addition, the pre-onset mean was removed from the post-onset signal (baseline correction). For each k epochs with m frequency bins between 15 and 35 Hz, the power spectral density (PSD) was computed in order to exclude poor trials. Following that, the Z-scored PSD error was calculated and those trials whose Z-scored PSD error was larger than 2 were omitted from the analysis (Average=1.2 with the range (0,5)). The mean (SD) number of trials that passed this quality control and were used for the hits analysis was 21.7 (3.6) for 1-back, 19.9 (3.6) for 2-back and 17.8 (3.8) for 3-back.For the non-target correct rejections, the mean (SD) number of trials that were used for further analysis was 45.7 (2.8) for 1-back, 43.3 (4.3) for 2-back and (5.6) for 3-back. Based on previous research, we selected the electrodes of the frontal (stimulated areas), central and parietal regions (Fz, F3,, F4, Cz and Pz channels) and the ERP components P200 (positive deviation with a peak of about 100-250 ms after stimulus presentation) and P300 (positive deviation with a peak of about 250-600 ms after stimulus presentation) for analysis [19,21]. In this study, the mean amplitude measure was used to analyze ERP components [27]. For the P200, mean amplitude of time window (100-250) was calculated [19], and for the P300, mean amplitude of time window (250-600) was computed [28]. All of these steps were performed in MATLAB 2020a.

##### Statistical analysis

For the statistical analysis, we computed the change between Post and Pre-interventions (Post-Pre) for P200 and P300 ERP in the yoga-sham tDCS and yoga-active tDCS interventions. The differences were determined by subtracting the Post-values from the Pre-values of P200 and P300 for each intervention across all individuals. To determine the influence of the intervention on changes in P300 and P200 values, a four-way within-subject repeated-measures ANOVA was conducted with intervention (active vs. sham), channel (Fz, F3, F4, Cz and Pz), working memory load (1-back, 2-back, 3-back), and trials (hits, non-target correct rejections) as the within-subject factors. Using boxplot techniques and the Shapiro-Wilk normality test, the normal distribution of data and the homogeneity of variances were examined. In the case of significant interaction and main effects (*P* <0.05), Bonferroni-corrected post-hoc paired t-tests were used to compare each pair of variables. Statistical analysis used the RStudio version 1.4.1103 (RStudio Inc., USA).

## Results

Among the 18 individuals who underwent analysis, 14 (77.78%) were single, 13 (72.22%) were male, and 16 (88.89%) were right-handed (Table 1). The mean (SD) age of the participants was 27.04 (4.51). The treatment was well tolerated by the subjects, and no significant adverse effects were seen during or after electrical stimulation. There was no difference in tDCS adverse impact estimates between interventions (Supplementary materials).

**Table 1:** Demographics.

|  |  |  |
| --- | --- | --- |
| **Sex, n (%)** | Male | 13 (72.22%) |
|  | Female | 5 (27.78%) |
| **Age, mean (SD)** |  | 27.04 (4.51) |
| **Marital status, n (%)** | Never Married | 14 (77.78%) |
|  | Married | 1 (5.56%) |
|  | Divorced | 1 (5.56%), |
|  | Domestic Partner | 2 (11.11%) |
| **Handedness, n (%)** | Left | 1 (5.56%) |
|  | Right | 16 (88.89%) |
|  | Ambidextrous | 1 (5.56%) |
| **Employment, n (%)** | Student | 5 (27.78%) |
|  | Working | 13 (72.22%) |

### Working memory-related ERP components

For the ERP components, significant interactions of intervention by channel by working memory load (*F* = 3.11, *P* = 0.001, η_p_^2^ = 0.024), intervention by channel (*F* = 2.87, *P* = 0.022, η_p_^2^ = 0.011), and intervention by working memory load (*F* = 3.78, *P* = 0.023, η_p_^2^ = 0.007) was found for the P200 component. The pairwise paired t-test comparison showed significant differences for changes of P200 for 2-back (p=0.048) in Fz electrode, P200 for 2-back (p=0.048) in F3 electrode among active and sham interventions for the hits in both electrodes, the P200 amplitude changes were increased in the active intervention compared to the sham.

In addition, significant interaction of intervention by working memory load (*F* =3.72, *P* = 0.024, η_p_^2^ = 0.007) was found for the P300 component. The pairwise paired t-test comparison showed significant differences for changes of P300 for 3-back (p=0.018) in F3 electrode for the hits and changes of P300 for 3-back (p=0.029) in Pz electrode for the non-target correct rejections. In addition, the findings revealed an increase in the P300 amplitude in the Pz and a reduction in the F3 channel P300 amplitude among active and sham intervention. We did not find any significant differences in the other interactions and electrodes. Figures 3 and 4 depict the grand averaged ERP plot (hits) of all subjects for pre-active, post-active, pre-sham and post-sham interventions for the target correct responses (hits), as well as the box plot of changes in P200 and P300 amplitude for the Fz and F3 channels. Figures 5 shows the grand averaged ERP plot (non-target correct rejections) of all subjects for all interventions for the non-target correct rejections, and the box plot of changes in P200 and P300 amplitude for the Pz channel.

**Figure 3:**
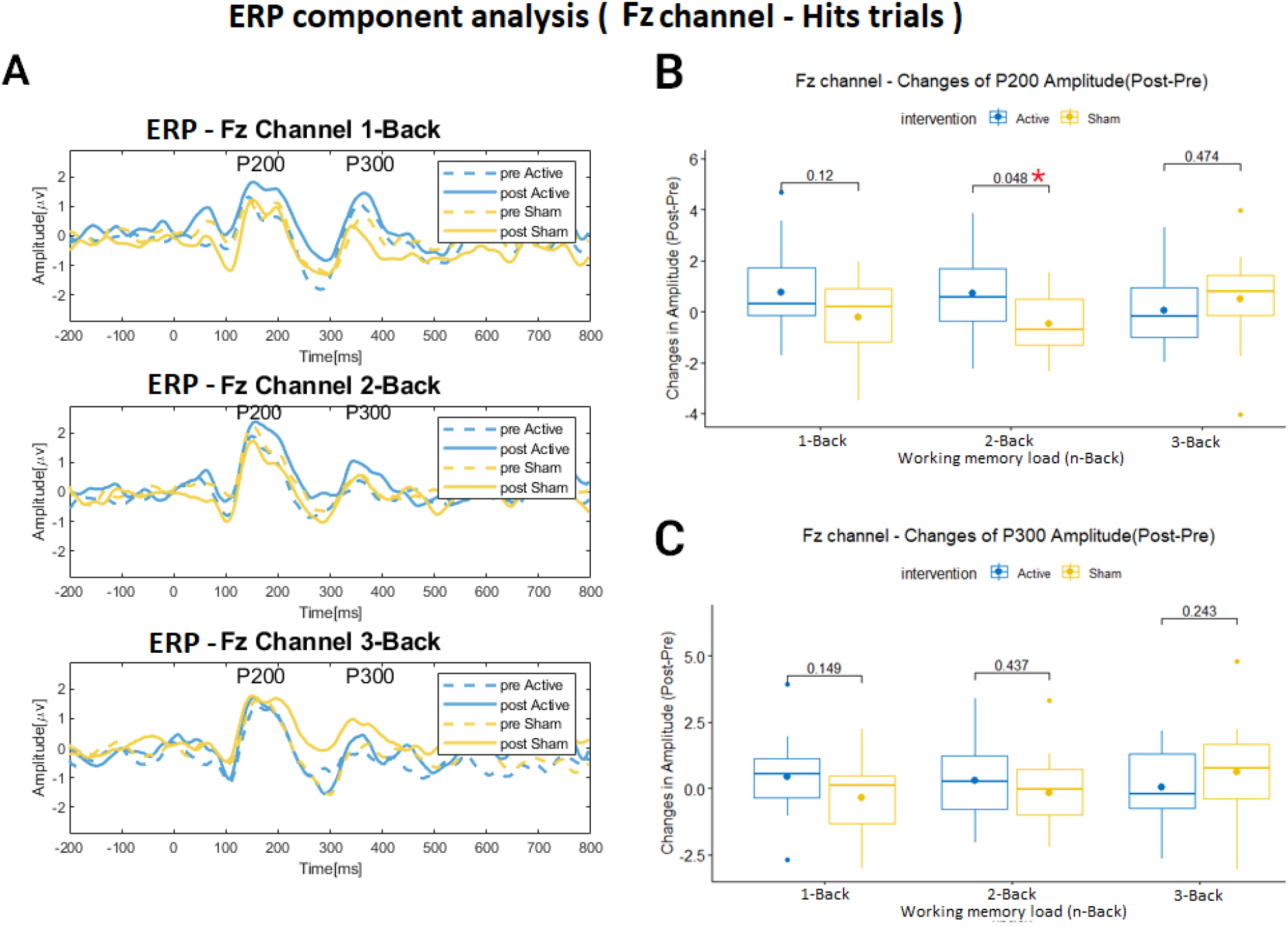
ERP component analysis (Fz channel, Hits trials) A – Grand averaged ERP plot(hits) for all subjects for pre-active, post-active, pre-sham and post-sham interventions for Fz electrode. The x-axis displays Time [ms], and the y-axis displays amplitude [µV]. B, C – The box plot of changes of P200 amplitude(B) and P300 amplitude(C) for different working memory loads (N-Back) for active (Post-Pre difference) and sham (Post-Pre difference) interventions for Fz electrode. The pairwise comparison showed significant differences for changes of P200 for 2-back(p=0.048). The box plot shows the median with a horizontal line, the mean with a dot, and the 25th and 75th percentiles with upper and lower boundaries. The whiskers show the 5-95 percentile. The x-axis shows working memory load (N-Back) and the y-axis shows changes in Amplitude[µV].

**Figure 4:**
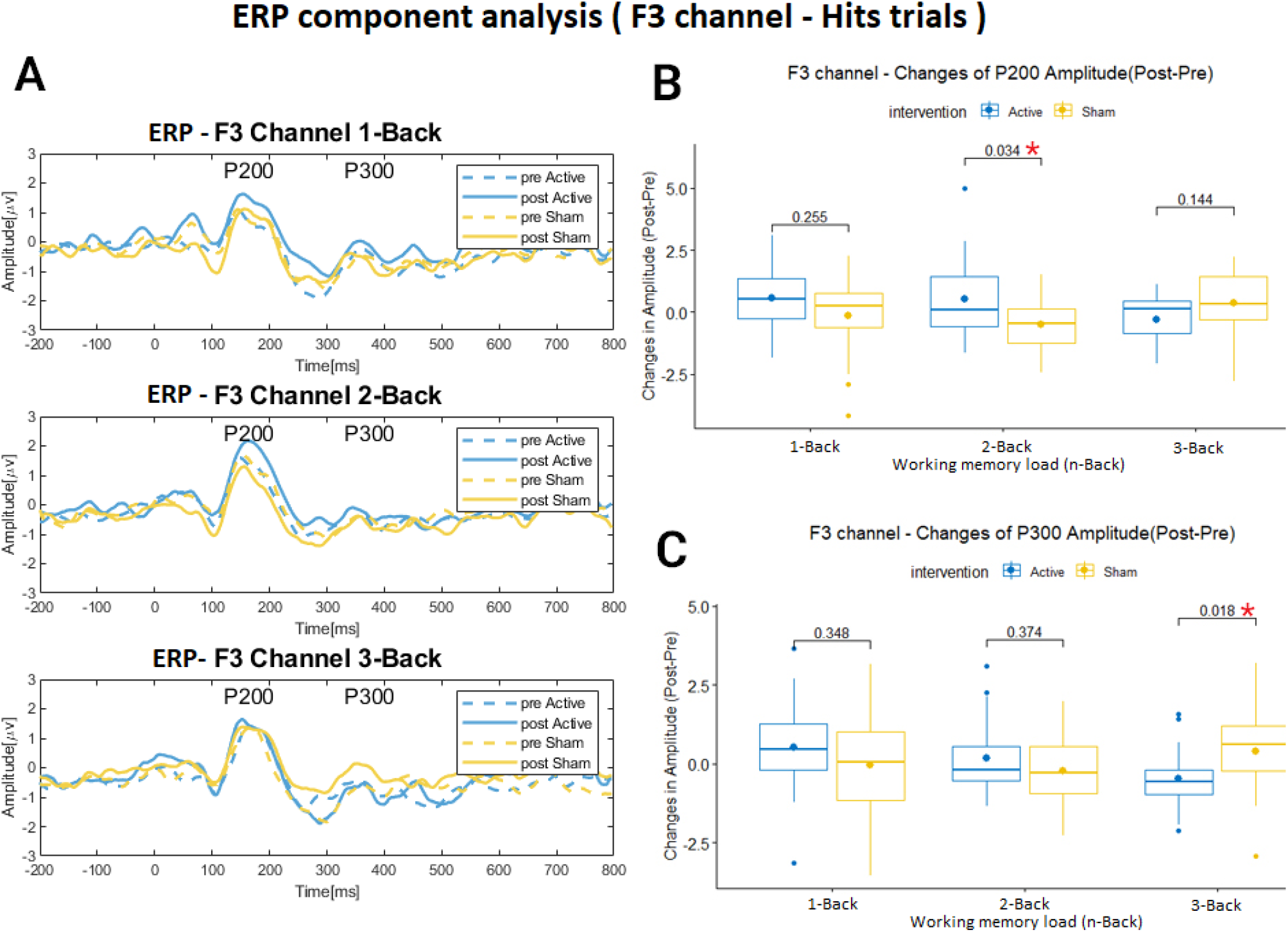
ERP component analysis (F3 channel, Hits trials) A – Grand averaged ERP plot(hits) for all subjects for pre-active, post-active, pre-sham and post-sham interventions for F3 electrode. The x-axis displays Time [ms], and the y-axis displays amplitude [µV]. B, C – The box plot of changes of P200 amplitude (B) and P300 amplitude(C) of different working memory loads (N-Back) for active (Post-Pre difference) and sham (Post-Pre difference) interventions for F3 electrode. The pairwise comparison showed significant differences for changes of P200 for 2-back(p=0.034) and changes of p300 for 3-back(p=0.018). The box plot shows the median with a horizontal line, the mean with a dot, and the 25th and 75th percentiles with upper and lower boundaries. The whiskers show the 5-95 percentile. The x-axis shows working memory load (N-Back) and the y-axis shows changes in Amplitude[µV].

**Figure 5:**
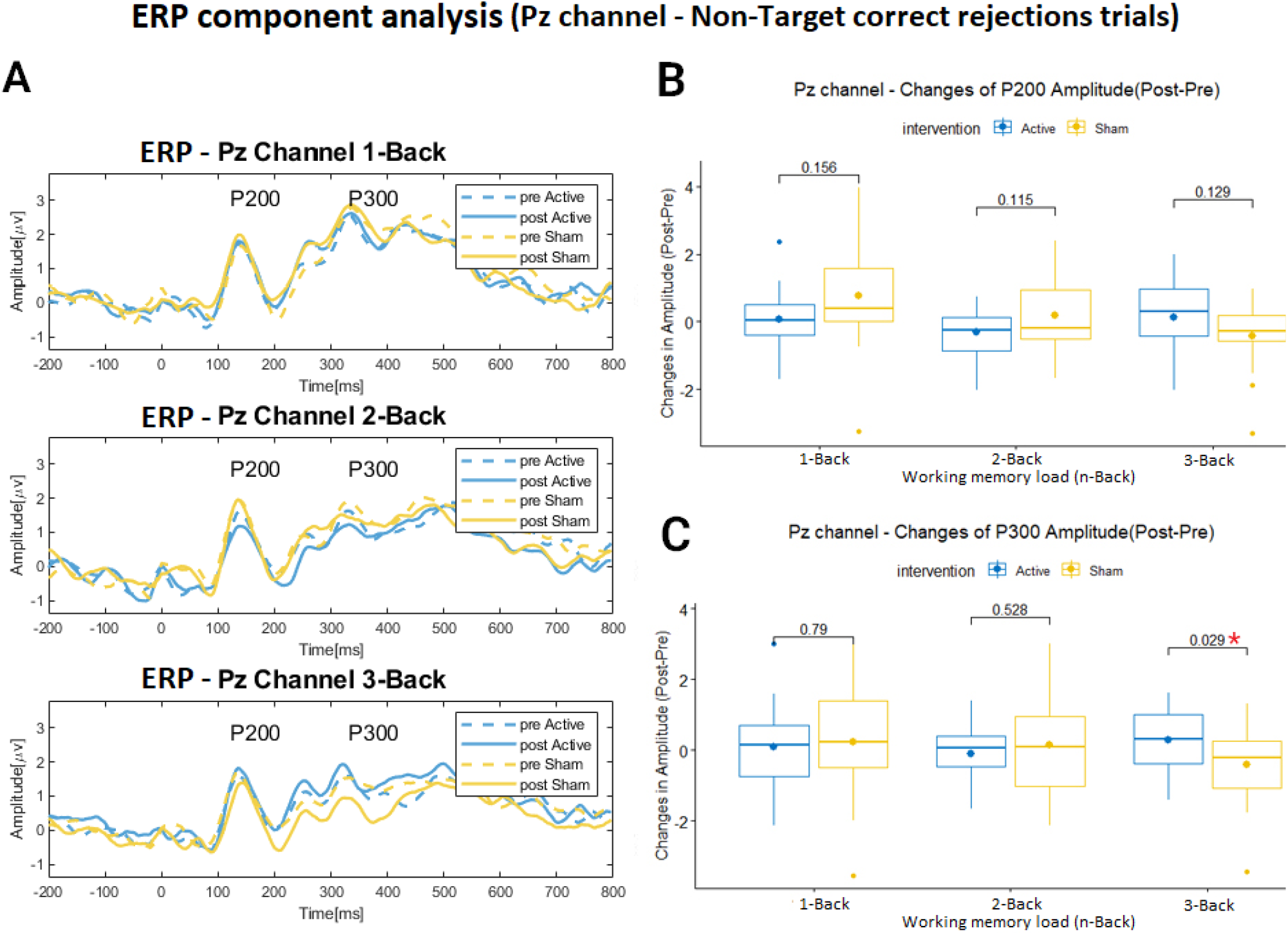
ERP component analysis (Pz channel, Non-Target correct rejections trials) A – Grand averaged ERP plot (non-target correct rejections) for all subjects for pre-active,post -active, pre-sham and post-sham interventions for F3 electrode. The x-axis displays Time [ms], and the y-axis displays amplitude [µV]. B, C – The box plot of changes of P200 amplitude (B) and P300 amplitutde(C) of different working memory loads (N-Back) for active (Post-Pre differrence) and sham (Post-Pre differrence) interventions for Pz electrode. The pairwise comparison showed significant differences for changes of P300 for 3-back(p=0.018). The box plot shows the median with a horizontal line, the mean with a dot, and the 25th and 75th percentiles with upper and lower boundaries. The whiskers show the 5-95 percentile. The x-axis shows working memory load (N-Back) and the y-axis shows changes in Amplitude[µV].

## Discussion

To our knowledge, this is the first study to investigate the combined impact of yoga and tDCS on ERP components elicited by n-back working memory tasks. Previous research has revealed that these methods separately influence neuronal activity and working memory [6,7,19]. The results of this study demonstrated significant differences in working memory neurophysiological signatures such as P200 amplitude during the 2-back task between combined yoga-active intervention and yoga-sham tDCS for the hits. These modifications are exhibited in the frontal electrodes Fz and F3 near the stimulation areas. Also, there is significant difference in the P300 changes in the F3 electrode for the hit’s trials and in the Pz electrode for the correct rejections at the 3-back level.

We observed significant changes in the P200 amplitude in the subjects that received yoga and active-tDCS compared to yoga and sham-tDCS for the hits. P200 spreads in the anterior-central and parietal-occipital regions of the brain, but its maximum amplitude is in the frontal area. It is evoked by visual stimuli and modulated by attention processes [28]. The P200, an attention-related component, is considered being the basis for subsequent cognitive processing by indexing early attentional recruitment and reflects perceptual processing [29], which is modulated by attention and elicited by visual stimuli [30]. The present study’s findings show significant changes in the P200 amplitude of the frontal electrodes, which is a component related to attention [28] and consistent with other studies [19]. Considering the effect of yoga exercises on attention [5] and simultaneously the impact of DLPFC tDCS on attention [31], current research on the combination of tDCS and yoga may change the attention process synergistically in the brain.

The changes in the P300 component between the two interventions at different electrodes and WM loads were significant in the F3 for 3-back(hits) and in the Pz for 3-back (correct rejections). Previous studies have shown that the changes in the P300 component are generally correlated with working memory capacities and behavioral results [32]. P300 shows a more robust distribution on the electrodes in the center of the wall on the scalp. Its amplitude increases with rare target stimuli, and its latency (at the peak of occurrence) changes with difficulty in detecting the target stimulus [28]. The findings of our previous studies with similar interventions have shown that the combining yoga and tDCS interventions have not led to a change in behavioral outcome related to working memory [33]. In the current investigation, EEG data were taken about thirty minutes following the completion of the yoga and tDCS interventions because to the cap setup time. This time delay may have led to the elimination of the intervention after effects. Another effective issue may have been the number of intervention sessions, which in this study was only one session. In another study, performing ten sessions of combining tDCS with meditation resulted in significant changes in working memory outcomes and the amplitude of the P300 on Fz and Pz [21]. Regarding to these limitations, which may affect the results, we found significant differences in P300 component for the 3-back in F3 and Pz electrode.

The previous study by our group [34] showed that yoga+tDCS was associated with connectivity changes between frontal areas and other parts of the brain (parietal) using resting state EEG. The current study also showed that the effects of the combination of yoga and tDCS interventions on ERP were accompanied by significant changes in neurophysiological signatures and frontal and parietal electrodes (F3, Fz, and Pz). These significant changes were in the frontal area for the hits (correct responses to the target stimuli) and the parietal area for the non-target correct rejections. Also, the results of this research indicated an increase in the P300 amplitude in the Pz electrode and a decrease in the P300 amplitude in the F3 in comparison of active and sham intervention for 3-back. These findings are in line with the neural efficiency concept, according to which increased cognitive ability is related to greater brain activity distribution (i.e., increase in parietal and decrease in frontal amplitudes) [21].

WM load differently affected ERP components in this research, which need to be considered. The results of this study found more significant changes in the 2-back working memory load for the P200 component in Fz and F3 electrodes than 3-back and 1-back for the hits. We find significant differences for the 3-back on F3 (hits) and Pz (correct rejections) electrodes and did not find significant changes in 1-back. It seems higher difficulty of WM to be affected by interjoined yoga and tDCS. The 2-back is the level of working memory, which was neither too easy nor too hard for the subjects. The 3-back requires more cognitive effort to answer and process the stimulus correctly for the subjects [28].

Our study had various limitations that should be noted. First, the limited sample size may have impeded the effectiveness of the statistical analyses to detect differences. Second, using the offline approach (EEG acquisition following the tDCS completion) and the time period between the conclusion of the intervention and the EEG data collection may have reduced the inferential power of the research. Finally, it is possible that anodal tDCS effects on brain excitability were later appearing, as shown in some works [35], which was not considered in this study. For future study, it is suggested to utilize alternate neuroimaging methods, such as fNIRS and fMRI, to evaluate intervention outcomes following some sessions, and to use various cognitive tasks for evaluation.

## Conclusion

In summary, our study showed that the combination of active tDCS and yoga was related to significant changes in the amplitude of the P200 component for the 2-back working memory task in Fz and F3 channels and P300 for 3-back in F3 electrode for the hits. Also, we found significant changes of the P300 amplitude for 3-back in Pz electrode for non-target correct rejections. These results imply that the combination of yoga and tDCS may result in neuronal alterations in the frontal (stimulation sites) and parietal areas associated with the n-back working memory task. These results suggest that combining behavioral and electrical neuromodulation techniques have the potential to enhance cognition and neurophysiological effects.

## References

1. Desai R, Tailor A, Bhatt T. Effects of yoga on brain waves and structural activation: A review. Complementary Therapies in Clinical Practice. 2015 May;21(2):112–8.

2. Ajjimaporn A, Rachiwong S, Siripornpanich V. Effects of 8 weeks of modified hatha yoga training on resting-state brain activity and the p300 ERP in patients with physical disability-related stress. J Phys Ther Sci. 2018;30(9):1187–92.

3. Kora P, Meenakshi K, Swaraja K, Rajani A, Raju MS. EEG based interpretation of human brain activity during yoga and meditation using machine learning: A systematic review. Complementary Therapies in Clinical Practice. 2021 May;43:101329.

4. Cahn BR, Polich J. Meditation states and traits: EEG, ERP, and neuroimaging studies. Psychological Bulletin. 2006;132(2):180–211.

5. Gothe NP, McAuley E. Yoga and Cognition. Psychosomatic Medicine. 2015 Sep;77(7):784–97.

6. Luu K, Hall PA. Hatha Yoga and Executive Function: A Systematic Review. The Journal of Alternative and Complementary Medicine. 2016 Feb;22(2):125–33.

7. Gothe NP, Khan I, Hayes J, Erlenbach E, Damoiseaux JS. Yoga Effects on Brain Health: A Systematic Review of the Current Literature. BPL. 2019 Dec 26;5(1):105–22.

8. Kang H, An SC, Kim NO, Sung M, Kang Y, Lee US, et al. Meditative Movement Affects Working Memory Related to Neural Activity in Adolescents: A Randomized Controlled Trial. Front Psychol. 2020 May 12;11:931.

9. Nitsche MA, Paulus W. Sustained excitability elevations induced by transcranial DC motor cortex stimulation in humans. Neurology. 2001 Nov 27;57(10):1899–901.

10. Ke Y, Wang N, Du J, Kong L, Liu S, Xu M, et al. The Effects of Transcranial Direct Current Stimulation (tDCS) on Working Memory Training in Healthy Young Adults. Front Hum Neurosci. 2019 Feb 1;13:19.

11. Mancuso LE, Ilieva IP, Hamilton RH, Farah MJ. Does Transcranial Direct Current Stimulation Improve Healthy Working Memory?: A Meta-analytic Review. Journal of Cognitive Neuroscience. 2016 Aug 1;28(8):1063–89.

12. Hill AT, Fitzgerald PB, Hoy KE. Effects of Anodal Transcranial Direct Current Stimulation on Working Memory: A Systematic Review and Meta-Analysis of Findings From Healthy and Neuropsychiatric Populations. Brain Stimulation. 2016 Mar;9(2):197–208.

13. Abellaneda-Pérez K, Vaqué-Alcázar L, Perellón-Alfonso R, Bargalló N, Kuo M, Pascual-Leone A, et al. Differential tDCS and tACS Effects on Working Memory-Related Neural Activity and Resting-State Connectivity. Front Neurosci. 2020 Jan 17;13:1440.

14. Baddeley A. Working Memory. Science. 1992 Jan 31;255(5044):556–9.

15. Eriksson J, Vogel EK, Lansner A, Bergström F, Nyberg L. Neurocognitive Architecture of Working Memory. Neuron. 2015 Oct;88(1):33–46.

16. Goldman-Rakic PS. Regional and cellular fractionation of working memory. Proc Natl Acad Sci USA. 1996 Nov 26;93(24):13473–80.

17. Miyake A, Shah P. editors. Models of Working Memory.Uk: Cambridge University Press; 1999.

18. Rugg M.D, Coles M.G. H. Electrophysiology of Mind. UK: Oxford University Press; 1996.

19. Keeser D, Padberg F, Reisinger E, Pogarell O, Kirsch V, Palm U, et al. Prefrontal direct current stimulation modulates resting EEG and event-related potentials in healthy subjects: A standardized low resolution tomography (sLORETA) study. NeuroImage. 2011 Mar;55(2):644–57.

20. Nikolin S, Martin D, Loo CK, Boonstra TW. Effects of TDCS dosage on working memory in healthy participants. Brain Stimulation. 2018 May;11(3):518–27.

21. Hunter MA, Lieberman G, Coffman BA, Trumbo MC, Armenta ML, Robinson CS, et al. Mindfulness-based training with transcranial direct current stimulation modulates neuronal resource allocation in working memory: A randomized pilot study with a nonequivalent control group. Heliyon. 2018 Jul;4(7):e00685.

22. Salehinejad MA, Wischnewski M, Ghanavati E, Mosayebi-Samani M, Kuo M, Nitsche MA. Cognitive functions and underlying parameters of human brain physiology are associated with chronotype. Nat Commun. 2021 Aug 3;12(1).

23. Imburgio MJ, Orr JM. Effects of prefrontal tDCS on executive function: Methodological considerations revealed by meta-analysis. Neuropsychologia. 2018 Aug;117:156–66.

24. Gandiga PC, Hummel FC, Cohen LG. Transcranial DC stimulation (tDCS): A tool for doubleblind sham-controlled clinical studies in brain stimulation. Clinical Neurophysiology. 2006 Apr;117(4):845–50.

25. Makoto’s preprocessing pipeline [Internet]. California San Diego :2022. [Accessed 20 March 2022]. Available from: https://sccn.ucsd.edu/wiki/Makoto’s_preprocessing_pipeline

26. Delorme A, Makeig S. EEGLAB: an open source toolbox for analysis of single-trial EEG dynamics including independent component analysis. Journal of Neuroscience Methods. 2004 Mar;134(1):9–21.

27. Nielsen K, Gonzalez R. Comparison of Common Amplitude Metrics in Event-Related Potential Analysis. Multivariate Behavioral Research. 2020 May 3;55(3):478–93.

28. Shalchy MA, Pergher V, Pahor A, Van Hulle MM, Seitz AR. N-Back Related ERPs Depend on Stimulus Type, Task Structure, Pre-processing, and Lab Factors. Front Hum Neurosci. 2020 Oct 28;14:549966.

29. Bourisly AK, Shuaib A. Neurophysiological effects of aging: A P200 ERP study. Translational Neuroscience. 2018 Jun 22;9(1):61–6.

30. Liu Y, Zhang D, Ma J, Li D, Yin H, Luo Y. The Attention Modulation on Timing: An Event-Related Potential Study. PLoS ONE. 2013 Jun 24;8(6):e66190.

31. Silva AF, Zortea M, Carvalho S, Leite J, Torres ILdS, Fregni F, et al. Anodal transcranial direct current stimulation over the left dorsolateral prefrontal cortex modulates attention and pain in fibromyalgia: randomized clinical trial. Sci Rep. 2017 Mar 9;7(1):135.

32. Scharinger C, Soutschek A, Schubert T, Gerjets P. When flanker meets the n-back: What EEG and pupil dilation data reveal about the interplay between the two central-executive working memory functions inhibition and updating. Psychophysiol. 2015 Oct;52(10):1293–304.

33. Danilewitz M, Gao S, Salehinejad MA, Ge R, Nitsche MA, Vila-Rodriguez F. Effect of combined yoga and transcranial direct current stimulation intervention on working memory and mindfulness. Journal of Integrative Neuroscience. 2021;20(2):367.

34. Sefat O, Salehinejad MA, Danilewitz M, Shalbaf R, Vila-Rodriguez F. Combined Yoga and Transcranial Direct Current Stimulation Increase Functional Connectivity and Synchronization in the Frontal Areas. Brain Topogr. 2022 Mar;35(2):207–18.

35. Agboada D, Mosayebi Samani M, Jamil A, Kuo M, Nitsche MA. Expanding the parameter space of anodal transcranial direct current stimulation of the primary motor cortex. Sci Rep. 2019 Dec 3;9(1):18185.

