## Supplementary Materials for "Enhancement of neurophysiological signatures of working memory by combined yoga and tDCS"

**Corresponding Author**

*Shared corresponding authors:

Reza Shalbaf

### CRQ Side Effects

Participants tolerated the stimulation well, with no serious adverse effects reported during and after electrical stimulation. No difference was found between ratings of tDCS side effects across interventions (Table S1).

**Table S1: CRQ Side Effects**

#### Active tDCS Intervention

| Side effects | mean | sd | min | max |
|---|---|---|---|---|
| Pain | 1.25 | 0.55 | 1 | 3 |
| Tingling | 2.85 | 1.81 | 1 | 6 |
| Burning | 1.70 | 1.30 | 1 | 6 |
| Fatigue | 1.85 | 1.60 | 1 | 7 |
| Nervousness | 1.16 | 0.50 | 1 | 3 |
| Concentration | 1.35 | 0.93 | 1 | 4 |
| Visual Perception | 1.00 | 0.00 | 1 | 1 |
| Headache | 1.00 | 0.00 | 1 | 1 |
| Uncomfortable | 1.36 | 0.50 | 1 | 2 |

#### Sham tDCS Intervention

| Side effects | mean | sd | min | max |
|---|---|---|---|---|
| Pain | 1.18 | 0.39 | 1 | 2 |
| Tingling | 3.82 | 1.97 | 1 | 9 |
| Burning | 1.95 | 1.99 | 1 | 9 |
| Fatigue | 2.05 | 1.28 | 1 | 5 |
| Nervousness | 1.18 | 0.39 | 1 | 2 |
| Concentration | 1.68 | 1.39 | 1 | 7 |
| Visual Perception | 1.23 | 0.43 | 1 | 2 |
| Headache | 1.05 | 0.21 | 1 | 2 |
| Uncomfortable | 1.47 | 0.52 | 1 | 2 |

### EEG Electrodes

Twenty-eight EEG electrodes from the scalp (Fp1, Fp2, F3, F4, F7, F8, Fz, C3, C4, Cz, FC1, FC2, FC5, FC6, P3, P4, Pz, CP1, CP2, CP5, Cp6, O1, O2, Iz, P7, P8, T7, T8) were selected in this study. The scalp region and electrodes have shown in figure S1.

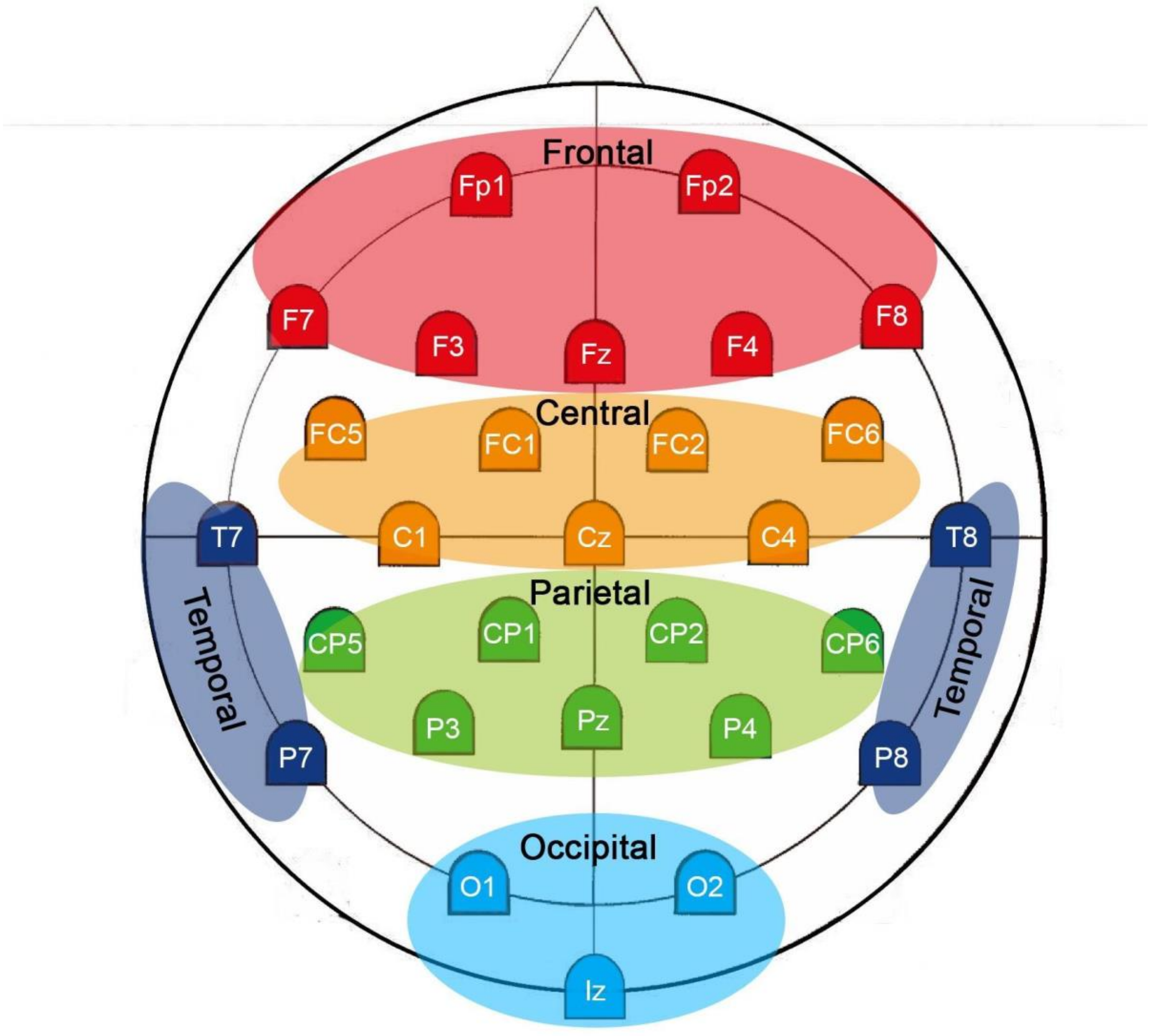


Figure S1: EEG Electrodes Region
